# Lateral pressure equalisation as a principle for designing support surfaces to prevent deep tissue pressure ulcers

**DOI:** 10.1101/592477

**Authors:** Colin J. Boyle, Diagarajen Carpanen, Thanyani Pandelani, Claire A. Higgins, Marc A. Masen, Spyros D. Masouros

## Abstract

When immobile or neuropathic patients are supported by beds or chairs, their soft tissues undergo deformations that can cause pressure ulcers. Current support surfaces that redistribute under-body pressures at vulnerable body sites have not succeeded in reducing pressure ulcer prevalence. Here we show that adding a supporting lateral pressure can counter-act the deformations induced by under-body pressure, and that this ‘pressure equalisation’ approach is a more effective way to reduce ulcer-inducing deformations than current approaches based on redistributing under-body pressure.

A finite element model of the seated pelvis predicts that applying a lateral pressure to the soft tissue reduces peak von Mises stress in the deep tissue by a factor of 2.4 relative to a standard cushion — a greater effect than that achieved by using a more conformable cushion. The ratio of peak lateral pressure to peak under-body pressure was shown to regulate deep tissue stress better than under-body pressure alone. By optimising the magnitude and position of lateral pressure, tissue deformations can be reduced to that induced when suspended in a fluid.

Our results explain the lack of efficacy in current support surfaces, and suggest a new approach to designing and evaluating support surfaces: ensuring sufficient lateral pressure is applied to counter-act under-body pressure.

## 1. Introduction

Supporting the body weight of critically ill, immobilised or paraplegic people without causing soft-tissue injury is not an easy task. The loading induced while lying or sitting for prolonged periods can cause damage to skin, adipose tissue and muscle; this damage is known as a pressure ulcer. Pressure ulcers are estimated to affect one in five hospitalised patients in Europe (1), while prevalence in some patient groups are much higher. For example, 85% of spinal cord injury patients develop a pressure ulcer over their lifetime (2) with associated care costs of approximately $1.2 billion annually in the US (3). A severe form of pressure ulcer develops in subdermal tissue close to bony prominences such as the ischial tuberosity and sacrum of the pelvis (4,5), and is known as a deep tissue injury. Because of the severity of deep tissue injury, preventative strategies have been a major focus in the field.

One approach to preventing pressure ulcers is to design support surfaces to reduce pathological pressures, and this has been a major area of research for the past forty years (6) - indeed ‘invalid beds’ have been developed since the 19^th^ century (7). While support surfaces have become increasingly high-tech, they have yet to outperform high-specification foam mattresses, and their adoption in clinics has not led to a significant reduction in pressure ulcer prevalence (8,9). While this lack of progress may indicate that we have reached the limit of support surface design, in this paper, we argue that current designs have been based upon a suboptimal design principle — that of under-body pressure re-distribution.

The presumption that high surface pressure leads to pressure ulcers, and therefore should be reduced, seems obvious. However, experimental, computational and clinical evidence suggests that high surface pressures do not necessarily cause pressure ulcers. Peak surface pressures (as measured by pressure mapping sensor arrays) cannot identify at-risk patients (10,11). High-tech mattresses that reduce peak surface pressures have increasingly been adopted in clinical settings yet their impact on pressure ulcer prevalence has been disappointing (8,9). Furthermore, soft tissue *can* tolerate extremely high surface pressures under certain circumstances. The soft tissues of a deep-sea diver, for example, are exposed to 100 kPa of surface pressure for every 10 m descended, yet pressure-related injuries to soft tissues are not a common issue in diving (12,13). Computational studies have helped to explain these observations, with Oomens et al. (14) demonstrating that peak surface pressure has very little impact on internal deformations near bony prominences — regions where deep tissue injuries are likely to occur (14,15). Since reducing peak surface pressure has failed to protect deep tissue, we sought to determine if there is any way to manipulate the external pressure profile that can protect deep tissue.

Deep tissue pressure ulcers develop as a result of several overlapping processes: ischaemia (16), ischaemia-reperfusion injury (17) lymphatic network obstruction (18,19) and direct cell deformation (20). Each of these processes is triggered by excessive deformation (exacerbated by shear stresses, microclimate, and other risk factors) within soft tissue, and so in some regards, a pressure ulcer could be more aptly named a ‘deformation ulcer’. Redistributing surface pressure (as current devices aim to do) does not necessarily reduce deformations (and hence pressure ulcers) because soft tissue has very different tolerances to the two components of stress (figure 1 a): deviatoric stress (which tends to change the shape of an object) and dilatational stress (which tends to change the volume of an object, but not its shape) (21,22). Human soft tissues are almost incompressible (23), and so can tolerate high dilatational stress with minimal deformation. In contrast, soft tissues deform readily with deviatoric stress, therefore it is this stress that must be minimised to prevent ulceration. While submersed, a diver experiences nearly uniform pressures on all surfaces; this tends to induce dilatational stress (22). On the other hand, interaction with very localised surface pressure – such as when sitting on a chair or lying on a mattress – induces large deviatoric stress (and therefore deformations) as the soft tissue bulges and is displaced laterally away from the load (figure 1 b).

**Figure 1.**
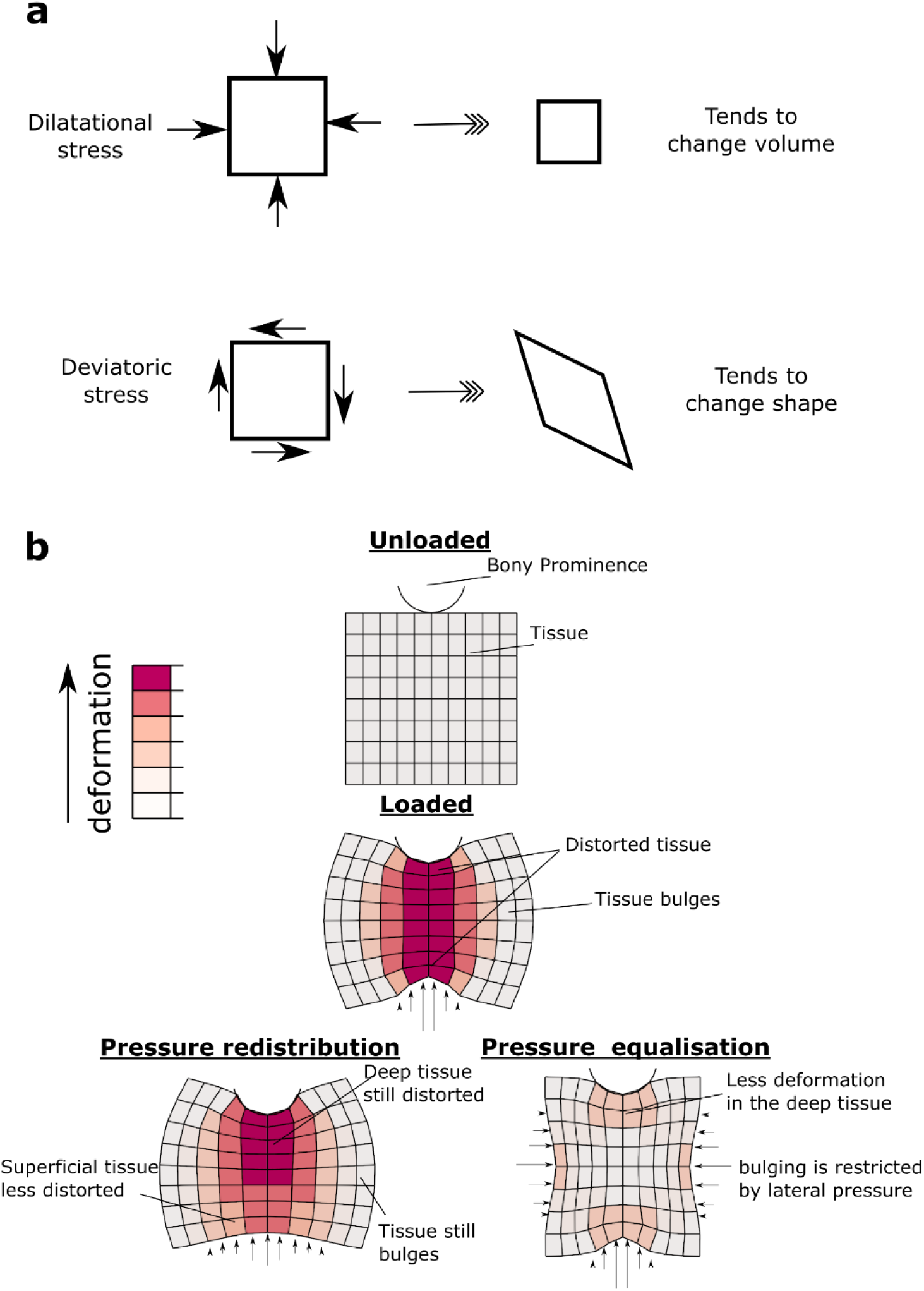
Deformations beneath a bony prominence. The stress in soft tissue has two components, dilatational and deviatoric (a). Soft tissue is much more resistant to dilatational stress than deviatoric stress. Under a bony prominence, the soft tissue is distorted due to the concentrated pressures at the bone and the support (b). Redistributing the surface pressure has some effect on the outer (superficial) region, but not on the deep tissue. We hypothesise that by applying pressure laterally (termed pressure equalisation), bulging is reduced, and the tissue can bear the load in a more dilatational mode.

One way to prevent excessive deformations may be to restrain the soft tissue from deforming by applying a supporting lateral pressure. In this paper, we test the plausibility of this principle using a computational model of the weight-bearing pelvis in a seated individual. We hypothesise that actively applying pressure laterally to the soft tissue of the pelvis will reduce the deformation at the ischial tuberosity to a greater extent than the commonly applied method of redistributing under-body pressure (figure 1b). Our rationale for this work is that by making subtle changes to the design objectives used for support surfaces, we may be able to substantially reduce the risk of ulceration for high-risk patients.

## 2. Methods

First, we adapted a previously-developed finite element model of seated buttocks (14) to test the hypothesis that applying lateral pressure will reduce tissue deformations (section 2.1). The seated position was chosen because the ischial tuberosity is a common site for deep tissue injury (1). Next, we used the model to determine whether applying lateral pressure or changing the stiffness of a standard cushion has the greatest effect on deep tissue deformations (section 2.2). To ensure that these effects translate to a more realistic setting, we developed a 3D model from MRI scans (section 2.3). Finally, we sought to formalise the relationship between surface pressure and internal deformations into a design principle — equalising under-body pressure with lateral pressure. To do this, we described the interaction of soft tissue and a support surface as a surface pressure boundary condition, which could be manipulated and studied independently of particular cushion design (section 2.4). All finite element input files and analysis protocols are supplied as supplementary data.

## 2.1 Model of the seated pelvis with lateral pressure application

### 2.1.1 Geometry and material models

An axisymmetric geometry was used to model the soft tissue surrounding a single ischial tuberosity in a seated individual. The geometry was similar to that used by Oomens et al. (14) but included more of the pelvis soft tissue to allow lateral pressure to be modelled (figure 2 a). The soft tissue was partitioned into fat, muscle and skin to produce similar patterns as found from MRI imaging (figure 2b). Each region was assigned an Ogden hyperelastic material model, and parameters were chosen to be consistent with the model described in Oomens et al. (14) (table 1). A flat, 76 mm-thick, two-layered seat cushion was modelled with hyperelastic material properties representing a soft cushion (table 1). An air-filled chamber (the pressure-equalisation device) was introduced to apply a lateral pressure to the soft tissues (figure 2 a). This was shaped to conform to the seated pelvis and was modelled using membrane elements that can resist tensile, but not bending, loads. The chamber wall material was modelled as sufficiently stiff (E = 10 MPa) so as not to allow appreciable changes in length.

**Figure 2.**
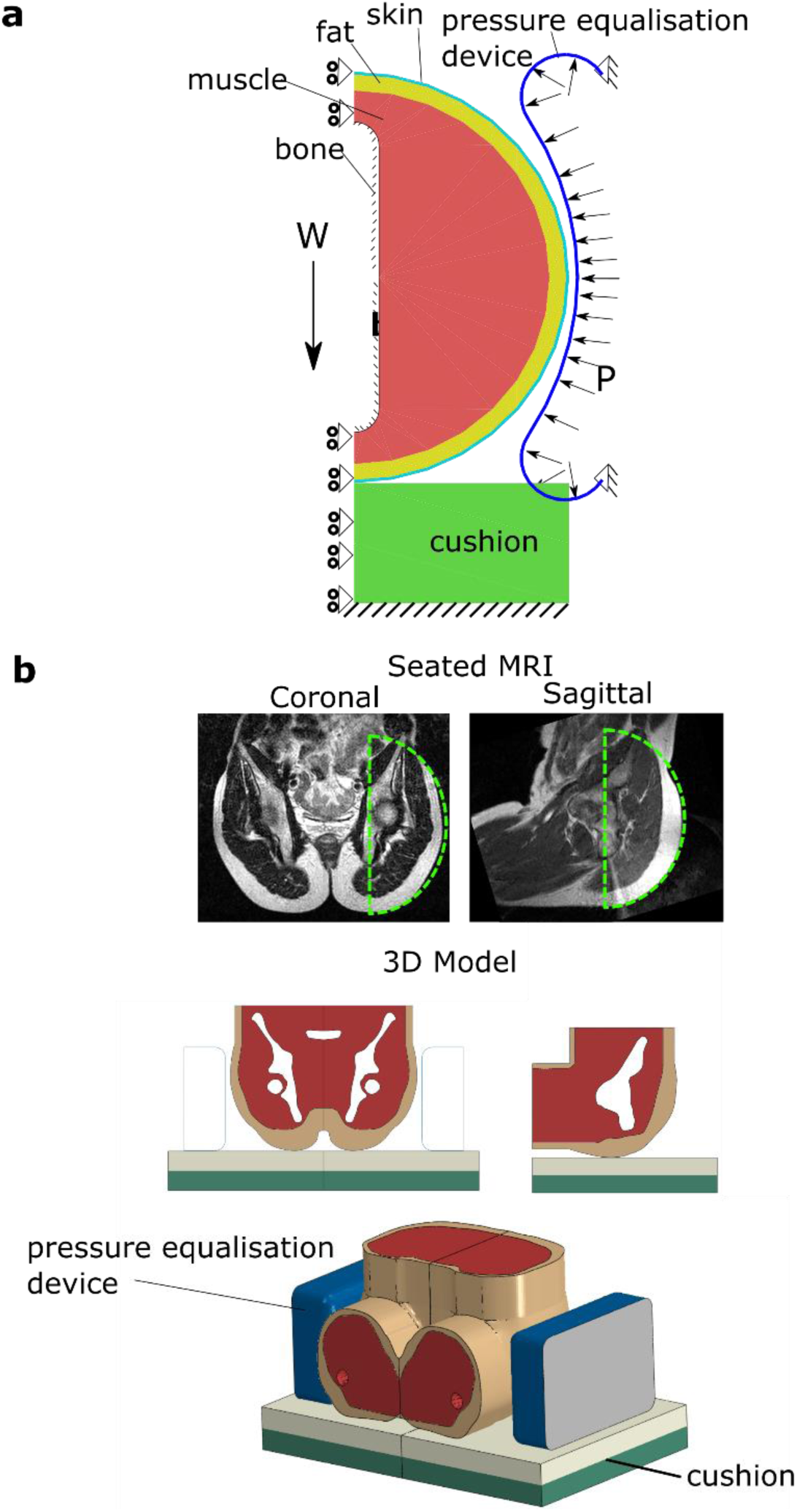
Finite element models. (a) An axisymmetric model of the soft tissue surrounding the ischial tuberosity. The model incorporates a rigid bony prominence, muscle, fat and skin layers interacting with a cushion and a pressure equalisation device. Axisymmetry was assumed, which allowed a force-controlled simulation of weight-bearing (W is the load borne by the ischial tuberosity). The pressure equalisation device was modelled as an air-filled chamber with a controllable internal pressure, P. (b) The axisymmetric region modelled is shown superimposed on saggital and coronal MR images of a seated male (top). A 3D model was generated from the MR images to assess 3D deformations (bottom).

**Table 1.** Parameters for the Ogden material model for each of the materials modelled. Ogden strain energy density function, 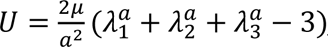, where *λ*_1,2,3_ are the principal stretches.

| Material | $\mu$ (MPa) | $a$ (-) |
| --- | --- | --- |
| Skin | 0.04 | 30 |
| Fat | 0.025 | 10 |
| Muscle | 0.045 | 5 |
| Stiff cushion | 0.08 | 10 |
| Medium cushion | 0.005 | 10 |
| Soft cushion top | 0.0035 | 7 |
| Soft cushion bottom | 0.005 | 10 |

### 2.1.2 Boundary Conditions

The amount of load supported by the pelvis was estimated at 400N — representing approximately 50% of the body weight of an 80 kg adult (14) — with each tuberosity bearing 200N. Symmetry boundary conditions were prescribed to all nodes lying along the z-axis (figure 2 a). The cushion base was constrained in all directions. Two nodes of the pressure equalisation device were constrained in all directions, and a uniform pressure was applied to the inner surface of the chamber to a maximum of 80. Frictionless contact was assumed between the support surfaces and the skin. Normal contact behaviour was enforced using the penalty method with finite-sliding (24).

### 2.1.3 Solution approach and output

A mesh sensitivity analysis was performed leading to a final mesh of 13056 linear quadrilateral elements representing the soft tissues. All models were solved as quasi-static, non-linear analyses using the ABAQUS finite element software v2016. To analyse and compare models, the von Mises stresses and shear strains in the soft tissue regions were calculated. von Mises stress is a scalar representing the deviatoric part of the stress tensor 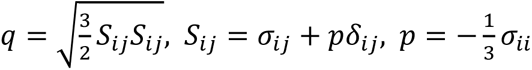. Shear strain was calculated as *ϵ*_1_ − *ϵ*_3_, where *ϵ*_1_, *ϵ*_3_ are the maximum and minimum principal strains. These were chosen to represent the level of deviatoric stress and strain, respectively.

Before summarising the stress and strain data, we adjusted to account for the varying volume of elements throughout the model. The data was re-sampled, with the probability of each point being chosen being the volume associated with that point (IVOL output from ABAQUS). Weighting the results like this ensures that analyses are independent of mesh density, which varies throughout the model.

Peak stress was defined as the 95th percentile of the stress data to avoid extreme outliers that may be sensitive to boundary conditions. Effects sizes (in the mean and peak values) between models were estimated by calculating bootstrapped 95% confidence intervals (25). We also analysed deep tissue stress along a path through the soft tissue directly beneath the ischial tuberosity, and along the surface of the ischial tuberosity.

### 2.2 Comparing lateral pressure application to changing cushion stiffness

Cushion stiffness is a design variable commonly used to create a more conformable cushion, and this was used as an intervention to compare to adding lateral pressure. The model described above was adapted to model load-bearing on three different cushion designs – a stiff, medium, and soft variety. The stiff and medium cushions were homogeneous, 38 mm thick cushions, while the most compliant (softest) was produced by defining a 76 mm thick two-layer cushion as in the previous section. The bottom layer of this cushion had the medium-stiffness cushion properties, and a softer material was assigned to the 38 mm top layer (table 1). These cushions were based on those analysed by Oomens et al. (14), which were in turn calibrated to model materials commonly used in wheelchair cushions.

For each of the three cushion simulations, load-bearing to 200N was established as in section 2.1. Pressure in the pressure equalisation device was then increased incrementally to a maximum of 80 kPa. The stresses and strains induced when lateral pressure is applied were computed.

### 2.3 Assessing deformations in 3D

Modelling the pelvis using a 2D axisymmetric model as above requires simplification of the bone and soft tissue geometries. To ensure that the beneficial effect of adding lateral pressure translates to 3D environments, we developed a 3D model that can more accurately capture the geometry of a seated pelvis. The MRI data of a male subject (age 30) was used to generate the 3D geometry (figure 2 b) including skin, fat, muscle and bone. Data usage was approved by the Imperial College Research Ethics committee under ethical approval number ICHTB HTA licence: 12275 and REC Wales approval: 12/WA/0196. To aid comparison of results between models, material properties for each layer were assigned to be consistent with the 2D model. The nodes representing the outer surface of the pelvic bones were constrained to move in the z-direction. Central symmetry was assumed to reduce model size by half. The soft tissue was meshed using 669,995 linear tetrahedral elements. A body force of 200N was applied to the bone nodes. This represents a full body weight of 80 kg, with the assumption that 50% of this travels through the pelvis of a seated individual (as was assumed with the 2D model and is based on Oomens et al. (14)). The lateral supports were modelled as air-filled cavities, with a thin outer membrane of material (as in the 2D model). To model the seated load case, the body force was ramped incrementally over the first analysis step. Next, the lateral supports were displaced towards the pelvis while the internal pressure within the support was fixed at 1 kPa. Finally, the pressure within the lateral support was increased to 10 kPa incrementally, and the von Mises stresses and maximum shear strains were calculated over ten increments. This model was solved using Abaqus/Explicit, with mass scaling applied to ensure kinetic energy was less than 1% of internal strain energy.

### 2.4 Determining the relationship between surface pressure and deep tissue mechanics

We studied how the shape and magnitude of the surface pressure distribution affected internal tissue stress to understand how these stresses can be minimised. The specific cushions and lateral support used in the previous sections were removed and replaced with a surface pressure boundary condition. For our axisymmetric model, pressure is a function of the angle from vertical, *P*(*θ*). The surface pressure was constrained to ensure that the model was in static equilibrium: the sum of the vertical forces due to surface pressure is equal to body weight, (*∮ P*(*θ*)cos*θds* = *W*), and the sum of the horizontal forces is zero (*∮ P*(*θ*)sin*θds* = 0).

The surface pressure on the buttocks when seated on a flat cushion follows a characteristic distribution (26) — there is a pressure peak beneath each ischial tuberosity which gradually reduces to zero towards the periphery of the contact area (supplementary figure 1). This contact pressure was modelled as a Gaussian distribution (see supplementary data). The spread of the pressure peak (*α*) was varied between 0.2 and 0.35, which represent cushions with stiffness values beyond the range of those tested in section 2.2. We then modelled an externally-applied lateral pressure by defining a second Gaussian term. This was controlled by its spread (*β*), its magnitude (*P_L_*), and its location (*θ*_0_). *β* and *θ*_0_ were fixed (0.4 and *π*/4 respectively) and *P_L_*/*P_V_* (the ratio of lateral pressure relative to under-body pressure) was varied from 0% to 75%. This led to 16 parameter combinations, described in table 2. Each pressure field was then applied as a boundary condition to the soft-tissue finite element model, and peak stresses and strains were calculated.

**Table 2.** Parameters governing the spread of the under-body pressure (*α*) and the magnitude of lateral pressure. Lateral pressure was defined relative to the peak under-body pressure (*P_L_*/*P_V_*).

| Pressure re-distribution ($\alpha$) | Lateral Pressure ($P_L/P_V$) |
| --- | --- |
| [0.5ex] 0.2 | 0 |
| 0.25 | 0 |
| 0.3 | 0 |
| 0.35 | 0 |
| 0.2 | 0.25 |
| 0.25 | 0.25 |
| 0.3 | 0.25 |
| 0.35 | 0.25 |
| 0.2 | 0.50 |
| 0.25 | 0.50 |
| 0.3 | 0.50 |
| 0.35 | 0.50 |
| 0.2 | 0.75 |
| 0.25 | 0.75 |
| 0.3 | 0.75 |
| 0.35 | 0.75 |

#### 2.4.1 Optimising the surface pressure distribution

We considered the pressure distribution of a body suspended in a fluid as an ideal support scenario (22), as it results in minimal deviatoric stress relative to dilatational stress (see supplementary data). We then optimised the location (*θ*_0_) and relative magnitude of the lateral pressure (*P_L_*/*P_V_*) to minimise the difference between *P*(*θ*) and the distribution when suspended in a fluid, while ensuring that the full body weight (200N) was supported (see supplementary data).

## 3. Results

### 3.1 Applying lateral pressure reduces soft tissue deformations

We first set out to determine the effect of adding lateral pressure to a person seated on a standard support surface. We used a finite element model of the pelvis to simulate weight bearing while sitting on a soft cushion. Firstly, we simulated weight bearing without lateral support. In agreement with other studies (14,22,27), the model predicts significant stress concentrations under the ischial tuberosity (figure 3 a), with peak von Mises stresses of 58 kPa produced in the muscle. Then, when lateral pressure is applied, the peak von Mises stress beneath the bony prominence drops to 18 kPa, in support of our hypothesis. Contour plots show the stress is more evenly distributed in the soft tissues (figure 3 a), suggesting that more of the soft tissue is being recruited in transferring the load.

**Figure 3.**
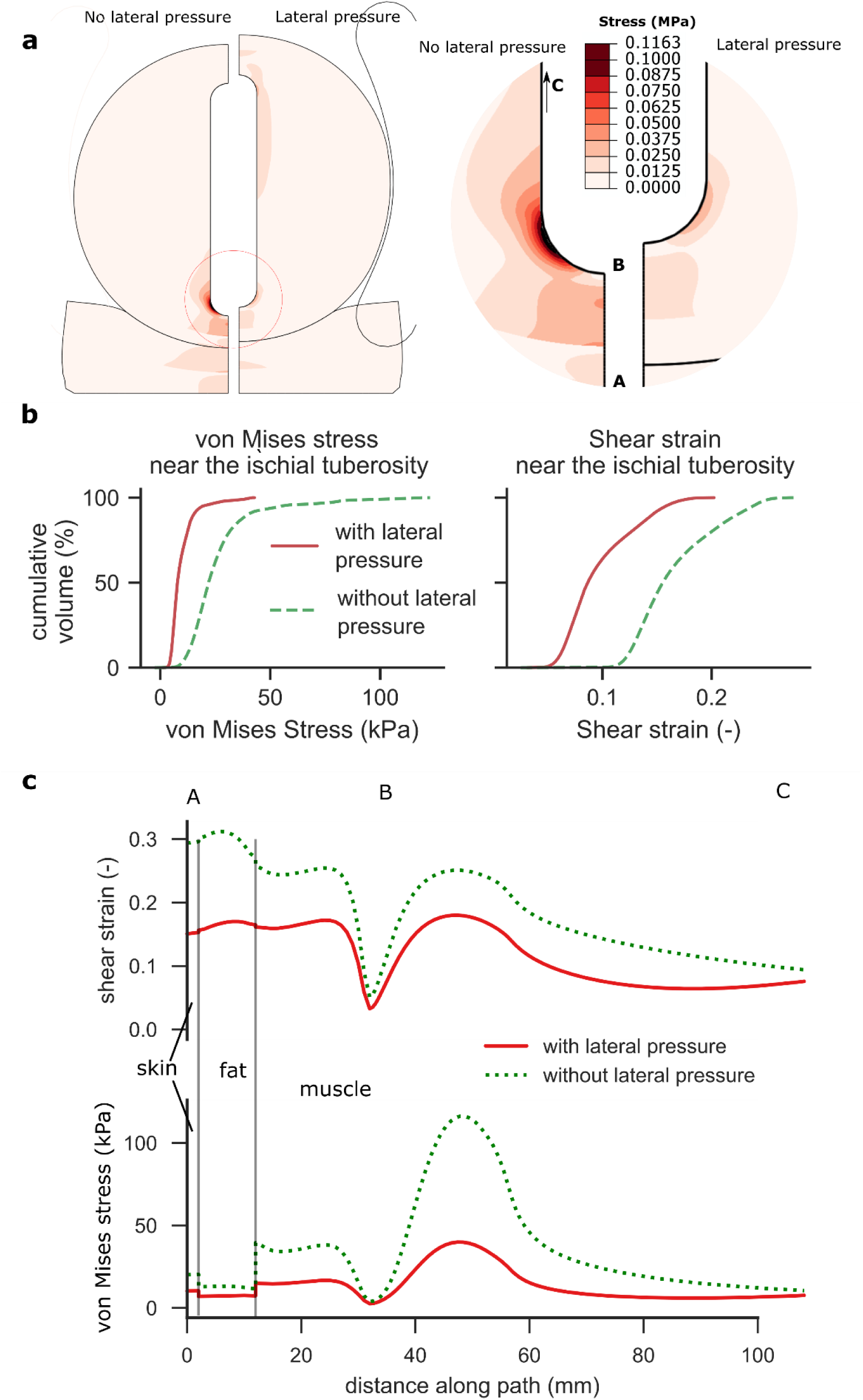
Analysis of load-bearing when seated on a soft cushion. In the absence of lateral pressure, the model predicts high von Mises stresses under the ischial tuberosity (a). With the introduction of lateral pressure (44 kPa chamber pressure), the region of high stress shrinks dramatically. Histograms of stresses and strains in the muscle tissue within a radius of 30 mm from the ischial tuberosity (b) indicate that von Mises stresses and shear strains are reduced. Analysis of the stress along path ABC (c) show a drop in von Mises stress and shear strain at the bony prominence, and throughout the muscle tissue. Shear strain and von Mises stress are also reduced in the skin and fat layers.

The volume of muscle tissue around the ischial tuberosity exposed to high von Mises stress (> 20 kPa) is reduced from 58% to 4% with lateral pressure application (figure 3 b). The volume of muscle tissue exposed to high shear strains (> 0.2) fell from 20% to 0%. Adding lateral pressure reduces the mean von Mises stress by 64% (95% CI 61% to 68%), while mean shear strains are reduced by 42% (95% CI 41% to 43%) when lateral pressure is applied (figure 3 b).

Plots of stress along a path through the soft tissue show that von Mises stress is reduced in all tissues under the ischial tuberosity when lateral pressure is applied (figure 3 c), with stress reduction being most pronounced in the muscle (66% in muscle, 43% in fat and 49% in skin). These results show that adding lateral pressure can reduce deep tissue deviatoric stress and deformation.

### 3.2 Applying lateral pressure reduces deformations to a greater extent than changing cushion stiffness

Having established that applying lateral pressure reduces deep tissue stress and deformation, we next assessed the effect of this intervention compared to a common device design consideration — changing cushion stiffness (figure 4). Cushion stiffness is usually manipulated to reduce peak pressures by maximising the contact area with the soft tissue. Indeed, the soft cushion provides approximately 3.5 times more contact area than the stiff cushion (72 cm^2^ in the stiff cushion, 177 cm^2^ in the medium cushion and 255 cm^2^ in the soft cushion), indicating that we are capturing a broad range of support surface stiffnesses. This area increase could be expected to achieve a similar reduction in deep tissue deformation. However, von Mises stresses (figure 4 a) and shear strains (figure 4 b) in the deep tissue are relatively less affected — with the change from a stiff to a soft cushion we see a reduction by a factor of 1.4 in peak von Mises stress (figure 4 c). Without lateral pressure, all cushions induce a peak von Mises stress > 50 kPa. We find that introducing lateral pressure reduces the peak von Mises stresses observed with each cushion by a factor of 2.4 on average (1.9 for the stiff cushion, 2.1 for the medium cushion and 3.2 for the soft cushion; figure 4 c). The lowest peak deep tissue stresses are observed when the soft cushion is combined with lateral pressure, which reduces the peak von Mises stress to 18 kPa, suggesting a synergistic effect of combining lateral pressure with a soft cushion (figure 4 c).

**Figure 4.**
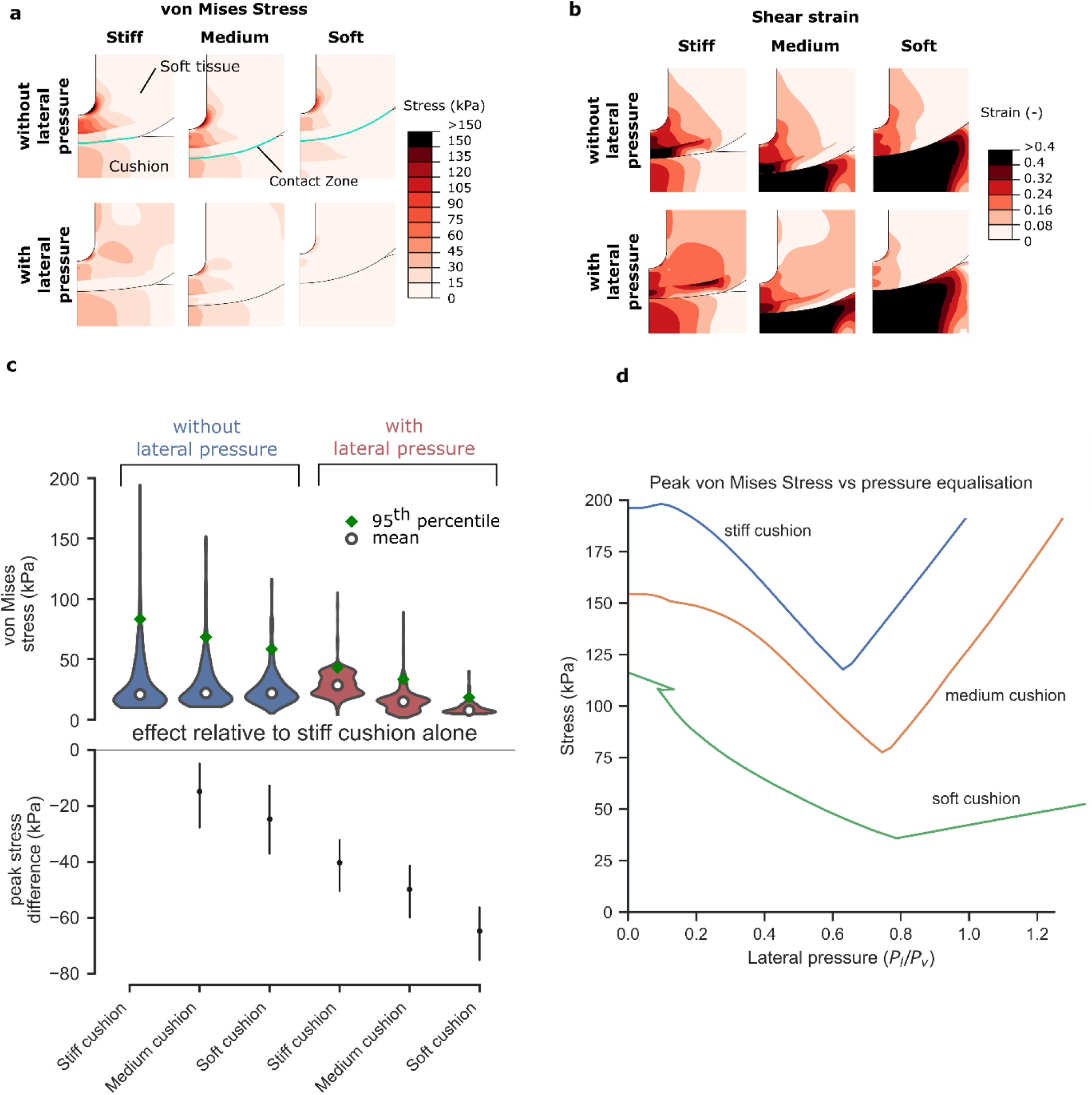
Applying lateral pressure is more effective than changing cushion stiffness. While the contact area varies substantially with cushion stiffness, the pattern of internal stress remains similar (a) — stress is concentrated at the bony prominence. Shear strains in the fat and skin are lower when a softer cushion is used (b), but strains within the muscle remain high for all cushions. These strains are reduced when lateral pressure is introduced. All three cushions benefit from the introduction of lateral pressure, with a soft cushion and lateral pressure providing the lowest von Mises stresses (c) [Violin plots show mean and 95th percentile values, stress difference plot shows the peak difference relative to a stiff cushion only with 95% confidence intervals]. As lateral pressure is gradually increased, the von Mises stress decreases until an optimum pressure is reached (d); beyond this pressure, von Mises stresses begin to increase again. While the magnitude of the optimum lateral pressure is different for each cushion, the ratio of lateral to vertical pressure is between 0.63 and 0.79 for all cushions tested.

We noticed that there is an optimum magnitude of lateral pressure which is different for each cushion (38.5 kPa, 37.9 kPa and 12.2 kPa for stiff, medium and soft cushions, respectively); however, the ratio of lateral to under-body pressure is consistently between 0.6 and 0.8 (figure 4 d). This suggests that balancing under-body and lateral pressures is more important for the reduction of deep tissue deviatoric stress than reducing peak under-body pressures.

### 3.3 Lateral pressure reduces stresses in a 3D model

Upon addition of lateral pressure, stresses were reduced at the ischial tuberosities in a similar way to the 2D model (figure 5 a). The volume of soft tissue exposed to high von Mises stress greatly reduced when lateral pressure was applied (figure 5 b). Adding lateral pressure reduced peak von Mises stress in the muscle by a factor of 2.5 when a stiff cushion was used, 2.6 with a medium-stiffness cushion and 2.4 with a soft cushion (figure 5 c). With optimal lateral pressure applied, the stresses at the greater trochanter and bones of the hemipelvis reached no more than 22% of the load at the ischial tuberosity. These results demonstrate that the effects found in 2D are representative of the 3D environment. They also show that gentle lateral pressure can be applied without compromising the tissue at the femur or sacrum. It should be noted that the lateral device modelled here was a very simple design, and no optimisation of its shape was performed. By contouring the lateral pressure device or other optimisations, further efficacy may be possible.

**Figure 5.**
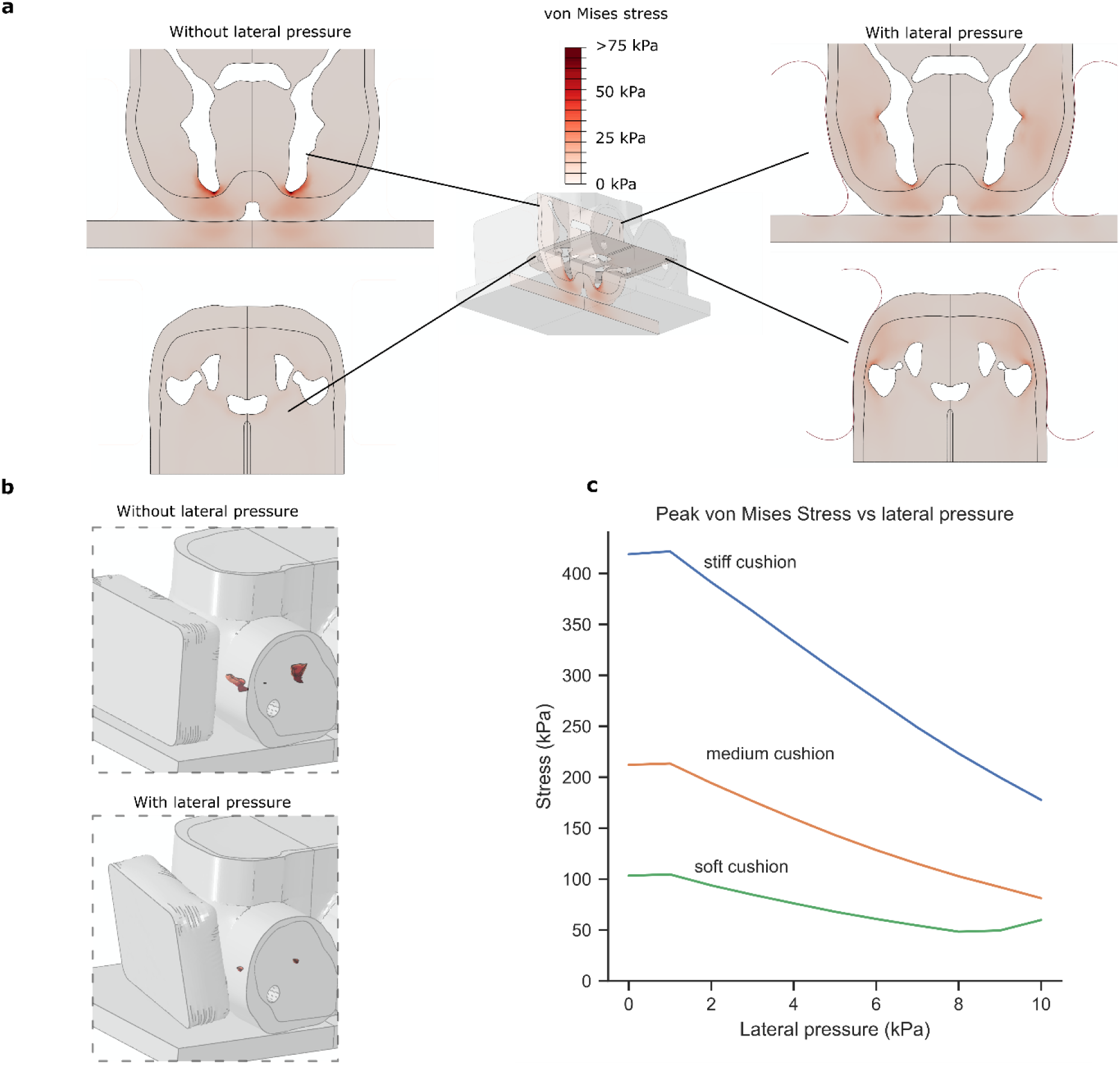
A 3D model of the seated pelvis under load. (a) Results for the stiff cushion shown with and without lateral pressure applied. Coronal and transverse sections are shown to indicate von Mises stresses both at the ischial tuberosities and the greater trochanter. (b) The volume of soft tissue exposed to high stresses (>32kPa) is shown in relation to the whole pelvis. The whole pelvis is made transparent to help visualise the location of high stresses (beneath the ischial tuberosity) (c) Change in peak von Mises stress throughout the soft tissue of the pelvis (surrounding both the ischium and the femur) as lateral pressure is increased.

### 3.4 Surface pressure equalisation is necessary to protect deep tissue from deformation

Having found that deep tissue deformations were minimised for each cushion when under-body and lateral pressure were in a specific ratio (0.6-0.8), we sought to test whether this ratio could form the basis of a design principle. We removed the cushion from the model and replaced it with a surface pressure boundary condition that could be manipulated independently of cushion design. We varied both the ratio of lateral to under-body pressure (*P_L_*/*P_V_*), and the spread of underbody pressure (*α*) and measured peak von Mises stress for all combinations (table 2).

As in the previous analyses, in the absence of lateral pressure, redistributing under-body pressure (increasing the spread, *α*, of the pressure peak), reduces peak von Mises stresses at the ischial tuberosity, but even substantial re-distribution fails to reduce the stress below 100 kPa (112 is observed when *α* = 0.35; figure 6 a). In contrast, inducing a lateral to under-body pressure ratio of 0.25 reduces peak von Mises stress from 180 kPa to 67 kPa. The presence of lateral pressure appears to reduce the effect of redistributing under-body pressure (figure 6 a), suggesting that when lateral pressure is employed, it becomes the most important factor in reducing deformations. These results indicate that controlling the ratio of lateral to under-body pressure (*P_L_*/*P_V_*) is necessary to achieve low deep-tissue stress.

**Figure 6.**
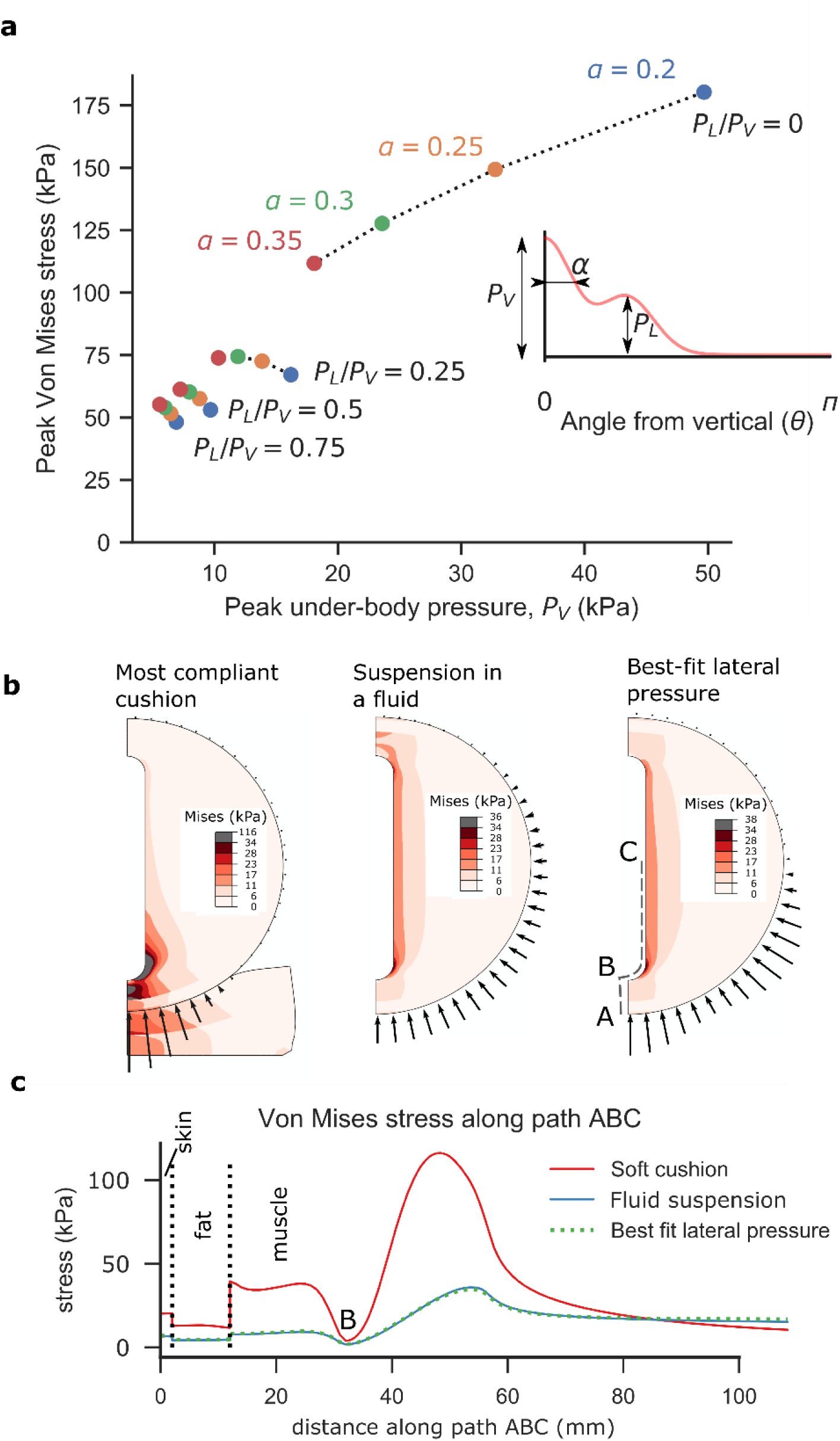
Surface pressure analysis. Redistributing under-body pressure (*P_V_*) reduces peak von Mises stresses when no lateral pressure is applied (a), but peak stresses remain above 100 kPa. Counter-acting that pressure with a lateral pressure (*P_L_*) reduces peak stresses to a greater extent. When the magnitude and angle of lateral pressure is optimised, the deep tissue von Mises stresses approach that of suspension in a fluid (b; arrows illustrate pressure intensity). Path plots of von Mises stress show that lateral pressure can induce a similar stress profile at the bony prominence to that when suspended in a fluid (c).

To understand why lateral pressure may be critical, we studied how this ratio affects the shape of the pressure distribution when compared to two extreme scenarios: the pressure distribution while sitting on a stiff cushion (a high-deformation scenario), and that when suspended in a fluid (a low-deformation scenario). The shape of these distributions are markedly different (supplementary figure 1), with a sharp peak of pressure beneath the ischial tuberosity when seated on a cushion, versus a smooth, even pressure distribution when submersed. A parametric study showed that adding lateral pressure best mimicked the pressure profile of suspension in a fluid (supplementary data).

We then optimised the magnitude and angle of the lateral pressure to best mimic suspension in a fluid. Contour plots show that optimising this lateral pressure (to *P_L_*/*P_V_* = 0.71 and *θ*_0_ = 61.5^∘^) can mimic the internal stresses experienced while suspended in a fluid (figure 6 b). In addition, wtih these parameters, von Mises stresses at the ischial tuberosity were either equal to or less than those induced when suspended in a fluid (figure 6 c).

In summary, not only can applying lateral pressure reduce deep tissue von Mises stress and deformation, we have found that an optimal magnitude and location of lateral pressure can mimic the environment induced when suspended in a fluid.

## 4. Discussion

The goal of reducing peak surface pressures at vulnerable body sites has underpinned the design of almost all medical support surfaces to date. Meanwhile, studies have consistently concluded that peak surface pressures do not accurately predict internal tissue mechanics (11,14), nor are they effective in predicting patients at risk of pressure ulcers (10). In this study, we have shown that ensuring under-body and lateral pressures are balanced — a principle we call pressure equalisation — is more effective at reducing deep tissue deformations than reducing peak under-body pressure. We postulate that devices designed to maintain a prescribed ratio of lateral pressure to under-body pressure will reduce the risk of pressure ulcer formation in the soft tissue of immobile patients.

The shift in emphasis from pressure re-distribution to pressure equalisation has implications for support surface design (figure 7). The synthesised results of multiple clinical trials (8,9) suggest that any well-designed mattress is better than a standard hospital bed, but none are particularly successful at reducing pressure ulcer risk. Pressure redistributing devices (either passive or active, figure 7 a and b respectively) may protect against superficial ulcers, while having little effect on deep tissue injuries (14). Our results show that devices must be capable of providing sufficient lateral support to counter-act the deformations induced by under-body pressure. Immersion/encapsulation-based devices such as water beds (28) aim to increase the contact area between the soft tissue and the support surface; however, while the contact area may increase, the horizontal pressures at the periphery of the contact area are usually minimal (figure 7 c), as pressure is primarily in reaction to gravitational body force. As this force acts in the vertical direction, it would be insufficient to equalise the under-body pressures and prevent bulging. Devices that aim to directly minimise shape change have also been developed (29), but their dependence on vertical translations of pistons means that they are not suited to regulating lateral pressure. We believe the advances in support surface technology may yet reduce pressure ulcer prevalence if they are redirected to achieving pressure equalisation.

**Figure 7.**
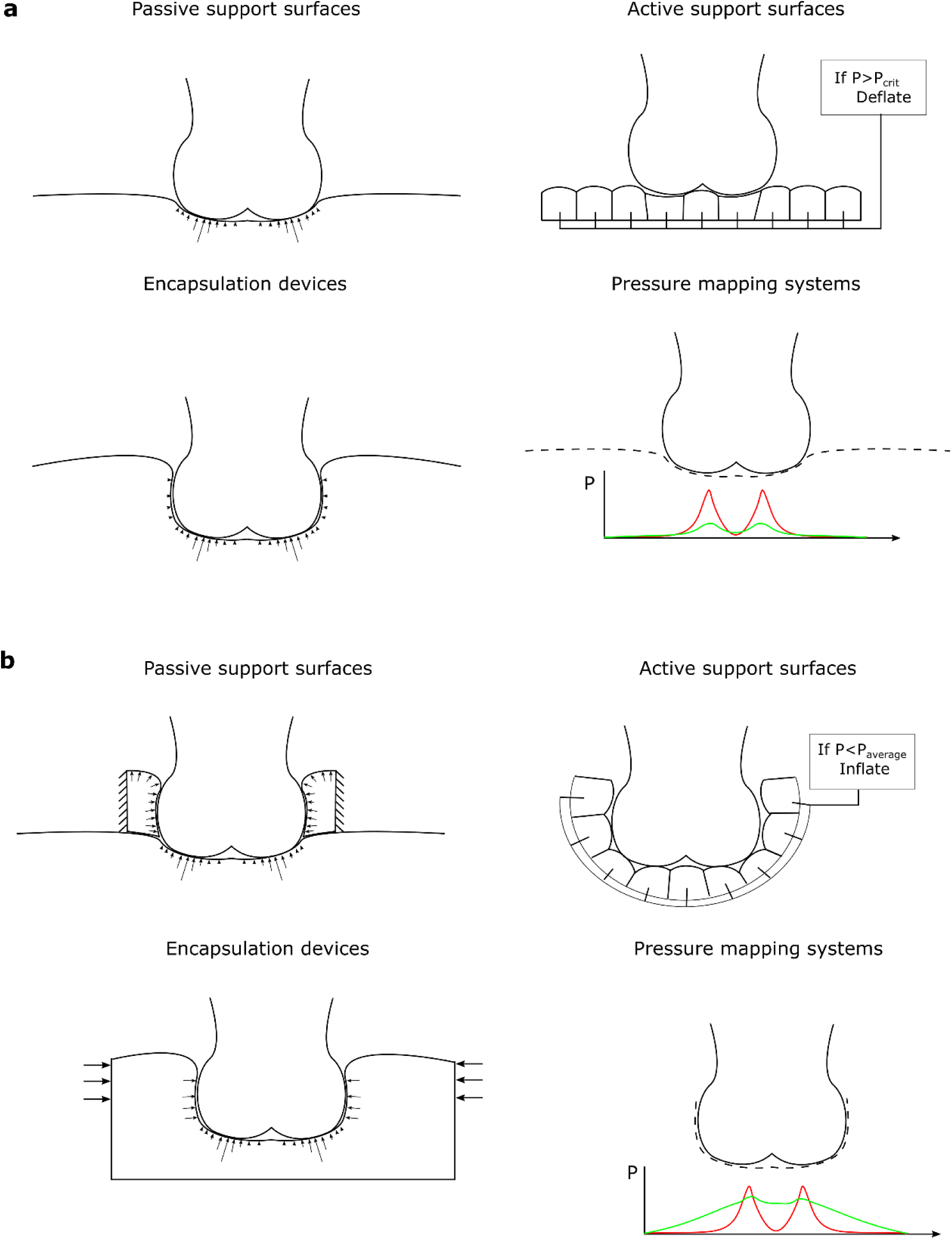
Pressure equalisation and its effects on device design. For surfaces designed to reduce peak pressure passively (a), applying a lateral pressure device helps to avoid lateral bulging (top, showing current devices, bottom showing improved design). Active devices based on individually controlled air cells (b) could be improved by surrounding the soft tissue, and changing the control software to aim for equalised pressure, rather than reduce peak pressure. Encapsulation devices achieve large contact areas, but the lateral pressures exerted may be limited (c). These could be improved by active compression or smart materials. Pressure mapping systems (d) currently identify pressure peaks as undesirable. If they could measure pressure around the surface, then they could be re-purposed to measure the level of pressure equalisation.

The pressure equalisation principle has implications for pressure measurement as a diagnostic tool and as a method of evaluating support surfaces. Surfaces incorporating arrays of pressure sensors have been suggested as early-warning systems for ulceration (30,31), and are frequently used to evaluate new support surfaces (32–34). Using this technology, devices can be readily differentiated based on the peak pressures they produce. However, when these devices are then compared through clinical outcomes, the differences between them vanish (8,9), and so the current predictive power of pressure measurement is limited. If surface pressure could be measured all around the soft tissues (figure 7 d), then the level of pressure equalisation may be a more predictive tool. Then, a measure of the ratio of lateral to under-body pressure (*P_L_*/*P_V_*) could be used to determine ulcer risk, and as a control signal for active devices.

The pressure gradient, defined as the spatial change in pressure from the point of peak pressure, has been proposed as an alternative to peak pressure for predicting soft tissue damage (35). At first glance, pressure equalisation may seem to be equivalent to using pressure gradient as an ulceration indicator. However, the pressure gradient does not account for the direction of pressure, and so a body could be loaded with a low pressure gradient yet have little or no pressure equalisation, because lateral pressures are not considered. In other words, pressure gradient is a local variable, as is peak pressure, whereas pressure equalisation is a measure of the quality of the body support as a whole.

From a clinical perspective, devices that can equalise under-body pressure with lateral pressure may be a vital tool in reducing the incidences of pressure ulcers — both hospital-acquired ulcers and those that are acquired in community care settings. The ability to apply sufficient lateral pressure will, however, need to be balanced with other equally important design considerations. For example, wheelchair users must not be exposed to intermittent high lateral pressures as they move in the chair. This inflexibility is one reason why form-fitted cushions are not a solution to preventing tissue distortion. In contrast, a successful device must be flexible enough to provide a well-distributed and well-controlled lateral pressure regardless of patient movement. Other design considerations include regulating the temperature and humidity at the skin surface, as well as ease of installation, use and cleaning. While this work has focused on the biomechanics of lateral pressure in general, the practical application of this principle will be more complex and require significant innovation in device design.

The reductions in deep tissue stress and strain possible through surface pressure equalisation could be sufficient to reduce pressure ulcer risk. The safe magnitude of deformation (and even the most appropriate measure) is not yet fully accounted for (36) and it is likely to be patient, environment, and tissue-specific. In this study, we have used two measures of deep tissue mechanics — von Mises stress and shear strain. These measures aim to capture the deformations likely to lead to capillary and lymphatic vessel restriction, and cell deformation, which contribute to pressure ulcer onset. Experiments using rat muscle under compression (37) indicated that stresses greater than 32 kPa induced damage, with this threshold dropping to 9 kPa over prolonged loading, while work quantifying deformations in a similar model (38) indicated that damage occurred above a shear strain of 0.3. Our results indicate that redistributing under-body pressure would not protect soft tissue from these levels of deformation, but that applying lateral pressure could.

The 2D finite element model used here simplified the anatomical structure of the pelvis in a similar way to previous studies (14,22,27). These idealisations allowed us to focus on the general case of a bony prominence transferring load through soft tissue to a support surface, and enabled the comparisons and analyses described here. The 3D model used here were important to support the conclusions drawn from 2D analyses, but there are also limitations to this model. While we have included more anatomical complexity including thighs, femurs and pelvis, more biofidelic models have been proposed (39–41). For example, we chose tissue mechanical properties in line with Oomens et al. (14), but there are several published models of soft tissue mechanical properties that vary in complexity. This makes conclusions based on absolute stress values difficult. For this reason, we have focused on the relative effects of interventions on stresses and strains, thus making the conclusions robust against the chosen material models. Using more biofidelic approaches will be a key step in applying the current results in the clinic. In particular, models generated from high-risk patients as opposed to healthy volunteers will be crucial. Physical validation will need to come from measurements of internal tissue deformations, for example through load-bearing MRI (42,43).

In conclusion, a change in focus from redistributing under-body pressure to equalising it with lateral pressure will lead to new innovations and improvements to patient care, resulting in a reduction of pressure ulcer prevalence in immobile patients.

## Data availability

All data required to reproduce the models in the paper are available at figshare. These include:

Abaqus input files for each of the cushion simulations, available at https://figshare.com/s/ddc34f0ae47835ef7a4b

Output data from the simulations, and Jupyter notebooks containing the data analysis used to generate the results and figures available at https://figshare.com/s/03727cdc9d7d76f9f216 for figures 3 and 4 and https://figshare.com/s/3a252a257dc0d2886b15 for figure 5 and supplementary data.

## Acknowledgements

We would like to thank Surbhi Gupta and all those involved at Imperial Innovations Ltd. for their support in delivering this project. This project was partly funded by an Imperial College London and Imperial Innovations Ltd. Proof of Concept funding to CB and SM and EPSRC funding to CH, MM and SM (EP/N026845/1).

## Supplementary Data

### S1 Analysis of pressure distributions

The surface pressure induced when seated on a cushion follows a characteristic shape (supplementary figure 1a): there is a narrow peak directly beneath the ischial tuberosity, occupying only a small region of the contact area. Pressure as a function of angle from the z-axis (supplementary figure 1 b) is adequately described using a gaussian function:

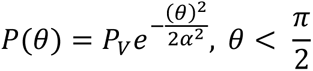

where *θ* is the swept angle between the vertical and the surface position, *α* controls the spread of the curve and *P_V_* is the peak pressure magnitude. Non-linear least squares fitting (using the curve_fit function of the scipy module) was used to fit the parameters for each of the three cushion types (table S1).

**Supplementary figure 1.**
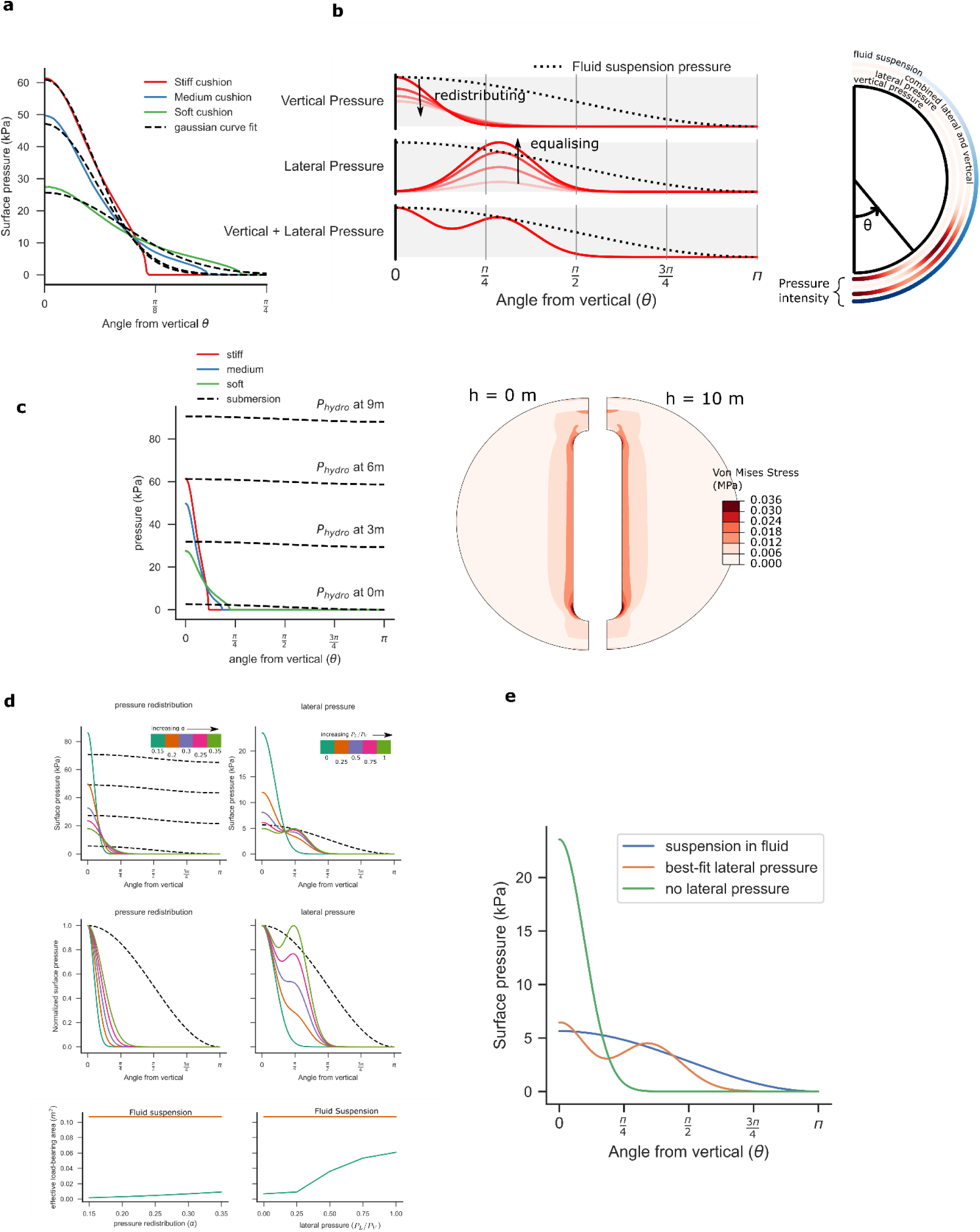
**a** Each of the cushions modelled in section 2.2 produces a similar pressure distribution at the skin surface: There is a peak directly beneath the ischial tuberosity (θ = 0), which drops to zero at the periphery of contact. Here, pressure is plotted as a function of the swept angle between the vertical and the position. A gaussian function provides a good fit for this distribution. **b** A schematic showing the lateral and under-body surface pressures. **c** The surface pressure induced when suspended in a fluid (dotted line) is relatively constant from each direction compared to the pressure profile when seated on a cushion (coloured lines). The magnitude of pressure varies substantially with depth, and even at 6 m below the surface, all surfaces are exposed to a pressure greater than that induced by a stiff cushion. When a pressure field representing 10 m submersion is compared to one representing pressure at the surface, the dilatational stress in the tissue increases (from 17kPa at 0m to 243kPa at 10m), but the deviatoric stresses are unchanged. **d** Both pressure re-distribution (increasing α) and applying lateral pressure (increasing P_L_/P_V_) reduces the peak pressure (top). Applying lateral pressure mimics the hydrostatic loading distribution more effectively than redistributing pressure (middle). The effective load-bearing area (area experiencing more than 50% of the peak pressure value) is increased more with lateral pressure than with re-distribution (bottom). **e** By optimising the magnitude and location of lateral pressure, the pressure profile around the soft tissue can accurately mimic that of suspension in a fluid (hydrostatic pressure).

**Table S1.** Parameters found through non-linear least square fitting of a gaussian distribution to the surface pressure.

| Cushion | $P_V(kPa)$ () | $\alpha$ (-) |
| --- | --- | --- |
| [0.5ex] Stiff | 60.9 | 0.170 |
| Medium | 47.1 | 0.184 |
| Soft | 25.7 | 0.272 |

### S2 Pressure distribution while suspended in a fluid

Suspension in a fluid can be regarded as a best-case scenario for load-bearing ^20^. In this case, the pressure distribution is given as:

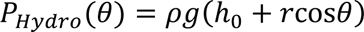

where *ρ* is density (997 for water), g = 9.81, *h*_0_ is the depth of submersion from the fluid surface to the model centre and *r* is the outer surface radius (130). The shape of the distribution remains constant as depth is increased, while the magnitudes of the pressures increase (supplemetary figure 1c). The dilatational stress within the soft tissue increases (from 17kPa when h0=130mm to 243kPa when h0=10130mm). In contrast, deviatoric stresses (von Mises) do not change at all as depth is increased.

### S3 Pressure re-distribution vs pressure equalisation

We modelled the introduction of a lateral pressure equalisation device by applying a new gaussian term, offset from vertical by angle *θ*_0_.

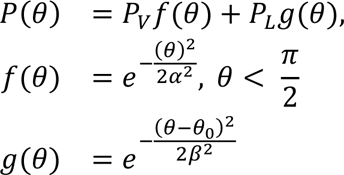

In these equations, *P_V_* and *P_L_* are the vertical and lateral pressure peak magnitudes, *α* and *β* control their spread, and *θ*_0_ controls the lateral peak location (supplementary figure 1b). Supplementary figure 1d shows that the ideal pressure profile (that of suspension in a fluid) is best achieved by applying lateral pressure. Taken from a different perspective, an objective of support surface design can be seen as trying to maximise the area of contact between the soft tissue and the cushion. Supplementary figure 1b shows that applying a small amount of lateral pressure increases the effective load bearing area^1^ far more than redistributing under-body pressure.

### S4 Scaling pressure distributions

Once the shape of *P*(*θ*) was described from any of the sections above, it was scaled such that it exerted a net force in the y-direction of 200 N, i.e. to support body force *W*. The force in the z direction acting on an infinitesimal area *dS* of a sphere of radius *r* is *dF_z_* = *P*(*θ*) ⋅ cos(*θ*)*dS*,

*dS* can be parameterised as *dS* = *r*^2^sin(*θ*)*dθϕ*, where *θ* is the polar angle and *ϕ* the azimuthal angle. The total force in the z direction is then:

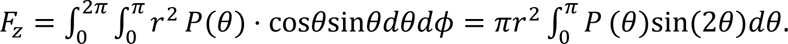

This was integrated numerically using Python scipy package (code included in figshare data).

### S5 Testing model assumptions

To test the effects of friction on model predictions, we set frictional contact in the 2D model described in the main paper (*μ* = 0.5). Analysis of von Mises stress predicted in models both with and without friction shows very small differences in peak stresses predicted (supplementary figure 2 a). Peak stress without friction and without lateral pressure applied was 116kPa, while with friction the maximum stress was 115kPa. Lateral pressure had a similar beneficial effect with and without friction (maximum stresses of 31kPa and 40kPa, respectively).

The tissue underlying the ischial tuberosity is patient-specific, and is either muscle or subcutaneous fat. In our 2D models, we assumed that muscle and subcutaneous fat was present. To test the effect of having just subcutanoeus tissue beneath the ischial tuberosity, we adapted our 2D model as shown in supplementary figure 2b. removing the muscle from under the ischial tuberosity resulted in a change in the location of the peak stress (to the overlying muscle, supplementary figure 2b). However, the beneficial effects of lateral pressure were still observed, with peak stresses reducing to 40kPa in both models.

**Supplementary figure 2.**
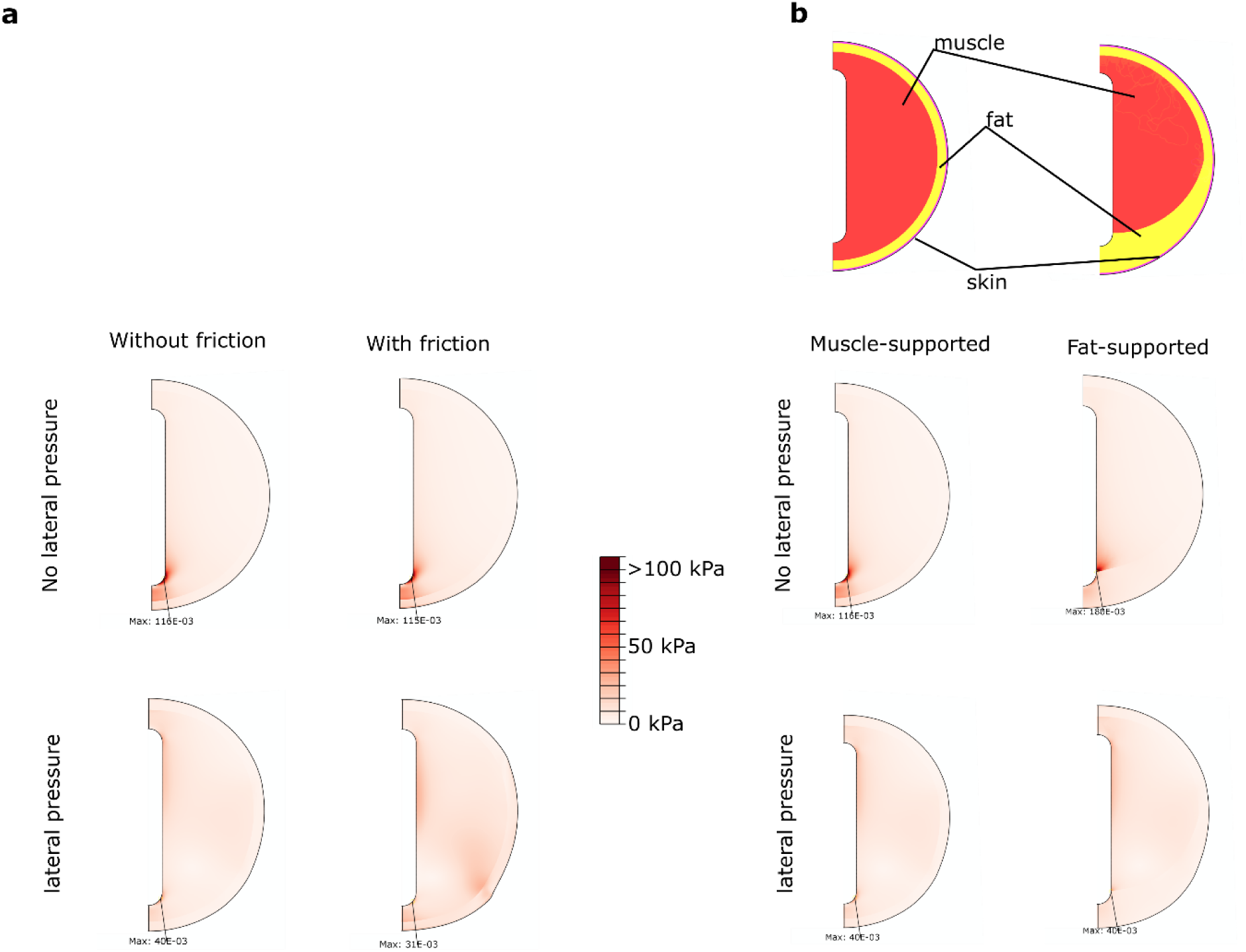
**a** von Mises stresses in models without (top) and with (bottom) lateral pressure. Original model with frictionless contact (left) versus the same mode with friction (right). **b** schematic showing the original model with underlying muscle tissue and an adapted model in which muscle does not lie under the bone. von Mises stresses shown without lateral pressure (middle) and with lateral pressure (bottom).

## Footnotes

1 This was defined as the surface area bearing at least half that of the peak pressure.

